# Modulation of cAMP metabolism for CFTR potentiation in human airway epithelial cells

**DOI:** 10.1101/2020.09.11.294223

**Authors:** Jenny P. Nguyen, Matthew Bianca, Ryan D. Huff, Nicholas Tiessen, Mark D. Inman, Jeremy A. Hirota

## Abstract

Cystic fibrosis (CF) is a genetic disease characterized by CF transmembrane regulator (CFTR) dysfunction. With over 2000 CFTR variants identified, in addition to known patient to patient variability, there is a need for personalized treatment. The discovery of CFTR modulators has shown efficacy in certain CF populations, however there are still CF populations without valid therapeutic options. With evidence suggesting that single drug therapeutics are insufficient for optimal management of CF disease, there has been an increased pursuit of combinatorial therapies. Our aim was to test cyclic AMP (cAMP) modulation, through ATP Binding Cassette Transporter C4 (ABCC4) and phosphodiesterase-4 (PDE-4) inhibition, as a potential add-on therapeutic to a clinically approved CFTR modulator, VX-770, as a method for increasing CFTR activity. Human airway epithelial cells (Calu-3) were used to test the efficacy of cAMP modulation by ABCC4 and PDE-4 inhibition through a series of concentration-response studies. Our results showed that cAMP modulation, in combination with VX-770, led to an increase in CFTR activity via an increase in sensitivity when compared to treatment of VX-770 alone. Our study suggests that cAMP modulation has potential to be pursued as an add-on therapy for the optimal management of CF disease.

## Introduction

Cystic fibrosis (CF) is a recessive genetic disease characterized by mutations in the gene encoding the CF transmembrane conductance regulator (CFTR), a phosphorylation-regulated ion channel responsible for conducting chloride and bicarbonate ions across epithelial cell membranes^[1–4]^. CFTR is localized at the apical membrane of epithelial cells that line primary conducting airways and submucosal glands and contributes to the regulation of airway surface lining (ASL) fluid volume and composition^[5]^. In CF, disease-causing mutations in CFTR lead to dysregulated ASL fluid volume and composition in the lung which is associated with increased mucus viscosity, increased susceptibility to pathogens, and modulated lung immunity^[6–9]^.

There are over 2000 known CFTR variants identified that are associated with a wide range of biological and functional consequences (www.genet.sickkids.on.ca). These variants have traditionally been classified into six classes based on their phenotype, including impairments to protein maturation, protein gating, and ion conductance^[10–12]^. However, evidence has suggested that this traditional classification is outdated since there exists CFTR variants with complex phenotypes that cannot be adequately classified into a single class^[13]^. Moreover, there has been evidence demonstrating that there are phenotypic differences between CF patients carrying the same genotypic CF variant^[14]^. This emphasizes the need for combinational therapies that will target several biological and functional defects.

Recent advancements in the molecular understanding of the functional impacts of specific CFTR variants has revolutionized drug development. As a result of discovery research focused on the function of specific CFTR variants, the Federal Drug Administration has approved four CFTR modulators^[15–18]^. The first CFTR modulator approved was CFTR potentiator ivacaftor (VX-770) which directly binds to CFTR to increase opening of the channel thus increasing chloride conductance^[19]^. In G551D-CFTR variants, VX-770 has demonstrated significant increases in predicted forced expiratory volume in 1 second (FEV_1_) from baseline at various doses and treatment durations^[20,21]^. VX-770 was subsequently assessed in combination with the CFTR corrector lumacaftor (VX-809), which increases the amount of CFTR reaching the cell surface, and showed mild improvements in FEV_1_ (2.6 - 4 percent points) and decreases in pulmonary exacerbations (30 - 39%) in the F508del homozygous population, which represents the global majority of CF patients^[22–24]^. Additional CFTR modulators approved include CFTR correctors tezacaftor (VX-661) and elexacaftor (VX-445)^[25–28]^. On-going combinatorial studies with VX-770, novel potentiators, VX-561 – which is an altered form of VX-770, ABBV-3067, and PTI-808; novel correctors, VX-661, VX-445, VX-659, VX-121, ABBV-2222, and PTI-801; and amplifier, PTI-428, which increases the amount of CFTR produced, suggest that a single drug therapy will not be sufficient for optimal management of CF disease (NCT03912233, NCT03969888, NCT03500263)^[25–30]^. Furthermore, although combinatorial CFTR modulator therapies have shown efficacy in previously poorly managed CF populations, many CFTR variants have not been studied at an individual level. In addition, these CFTR modulator therapies directly target CFTR defects and do not take into account mechanisms implicated in CFTR activity, such as phosphorylation of the regulatory domain of CFTR^[31–33]^. As such, new drugs and drug combinations that align with precision medicine approaches should be developed to ensure that there are valid therapeutic options for each individual with CF.

While understanding the genetic variation in CF patients is important for developing combinatorial therapies, there are many factors that can influence CFTR potentiation, including cyclic AMP (cAMP) metabolism. CFTR is phosphorylated by the cAMP-dependent enzyme protein kinase A (PKA)^[31,32]^. Following a rise in intracellular cAMP, PKA is activated and can phosphorylate CFTR, increasing open channel probability^[31–33]^. There are multiple mechanisms contributing to cAMP regulation, including cAMP-efflux transporters, phosphodiesterases (PDE), and ß_2_ adrenergic receptors^[14,34–39]^. A key cAMP transporter is ATP Binding Cassette Transporter C4 (ABCC4), formerly known as multi-drug resistance protein-4^[34,40]^. ABCC4 is functionally and physically coupled with CFTR through the scaffolding protein PDZK1^[34]^. Inhibition of ABCC4 with MK-571 is able to attenuate cAMP transport and potentiate CFTR activity in epithelial cells^[14,34,41,42]^. In the context of CF, we have demonstrated that pharmacological inhibition of ABCC4 in human airway epithelial cells is able to potentiate CFTR in G551D-variants above and beyond VX-770^[14]^. Aside from ABCC4 inhibition, there has also been evidence demonstrating that pharmacological inhibition of PDE-4 was able to potentiate CFTR activity through the elevation of cAMP levels^[37,38,43]^. The interaction of ABCC4 and PDE-4 inhibition to increase cAMP levels to augment CFTR potentiation remains to be explored.

Currently there are several investigational or clinically available therapeutics that have potential applications in influencing CFTR activity through cAMP elevation. MK-571 has been previously used as an ABCC4 inhibitor to prevent cAMP-efflux, resulting in an increase in CFTR activity in human airway epithelial cells^[14,41,42]^. MK-571 was designed as a cysteinyl leukotriene receptor antagonist and is exploited as a research compound due to off-target effects that include ABCC4 and PDE-4 inhibition and has also been used in humans safely^[44]^. Conversely, Ceefourin-1 and 2 have been reported as ABCC4 inhibitors with demonstrated *in vitro* efficacy and selectivity^[45]^. Roflumilast is a clinically available selective PDE-4 inhibitor with antiinflammatory effects that is commonly administered to chronic obstructive pulmonary disease (COPD) patients suffering from chronic bronchitis^[46–49]^. Similar to Roflumilast, Rolipram is also known to be a selective PDE-4 inhibitor. Originally it was investigated as a potential antidepressant drug, however, it is not used clinically due to adverse side-effects and its small therapeutic window^[50–52]^.

Due to our previous demonstration that pharmacological inhibition of ABCC4, in the presence of CFTR modulator VX-770, leads to CFTR potentiation beyond VX-770 alone and evidence demonstrating the use of PDE-4 inhibitors is also able to potentiate CFTR activity, we hypothesize that cAMP modulation with ABCC4 and PDE-4 inhibitors, in the presence of VX770, will potentiate CFTR activity. In order to begin defining the efficacy of cAMP modulation by ABCC4 and PDE-4 inhibitors on CFTR activity, we pursued a series of concentration-response studies on Calu-3 human airway epithelial cells with wild-type CFTR^[53–55]^. We demonstrate that combinatorial additions of CFTR modulator VX-770 with either an ABCC4 or PDE-4 inhibitor led to CFTR potentiation via an increase in sensitivity. Our results suggest that combinatorial additions of VX-770 with ABCC4 or PDE-4 inhibitors may increase sensitivity and efficacy of VX-770 alone.

## Results

### *In vitro* cAMP-efflux analysis of ABCC4 inhibitor compounds in human airway epithelial cells

An *in vitro* cAMP-efflux assay of two commercially available ABCC4 inhibitors, MK-571 and Ceefourin-1 (**Fig. 1a and b**), was performed in the human bronchial epithelial (HBEC6-KT) cells to define their efficacy in inhibiting cAMP-efflux. Concentration-dependent ABCC4 inhibition using MK-571 and Ceefourin-1 (**Fig. 1**) attenuation of extracellular transport of cAMP was observed. The half maximal inhibitory concentration (IC_50_) values of MK-571 and Ceefourin-1 were found to be 0.2 μM and 4.8 μM respectively and defined a role for ABCC4-mediated cAMP transport in human airway epithelial cells that may be leveraged for CFTR potentiation.

**Figure 1:**
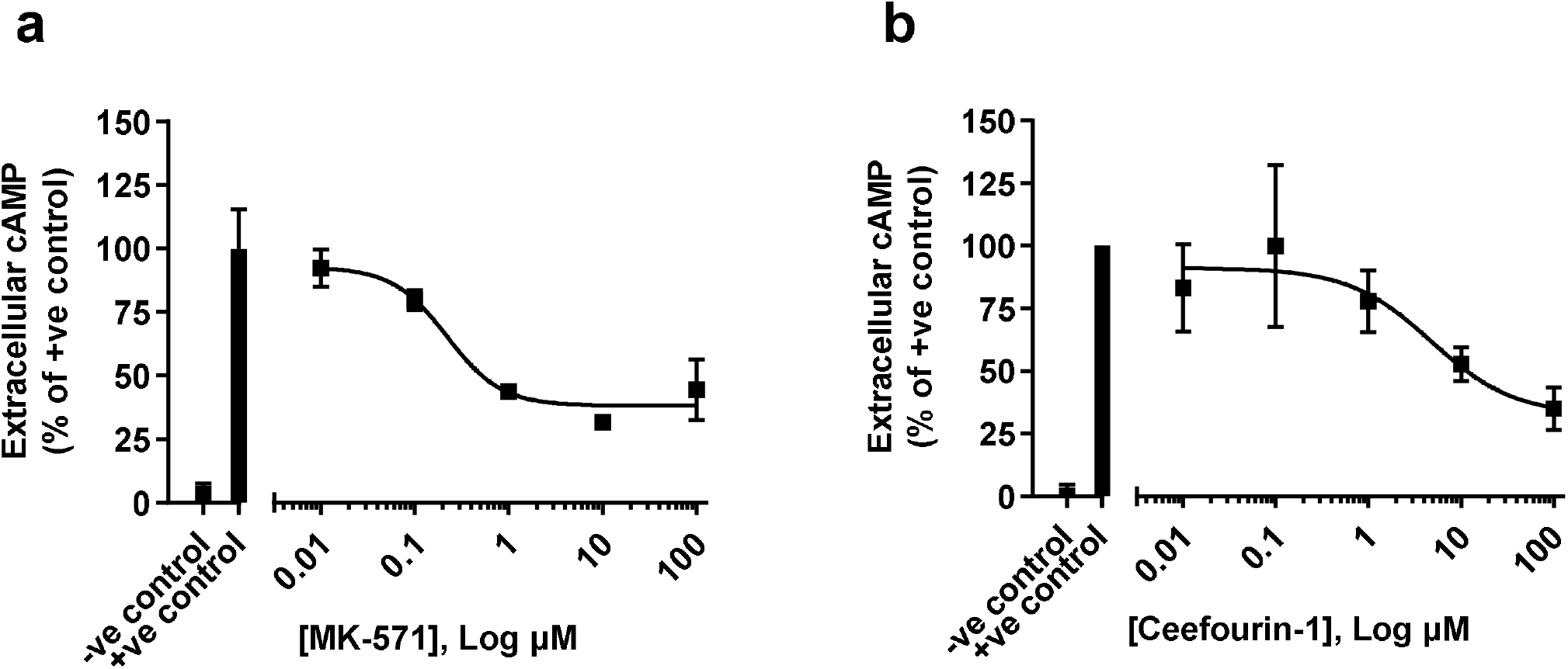
Concentration-response analysis of compounds with ABCC4 inhibition properties on cAMP-efflux from human airway epithelial cells. Human airway epithelial (HBEC6-KT) cells were pre-treated with IBMX (20 μM), exposed to (**a**) MK-571, (**b**) Ceefourin-1, or DMSO vehicle control, and then treated with forskolin (10 μM). Cell-culture supernatants were assessed for cAMP levels 24h post-treatment. Each concentration-response curve was normalized to positive control (IBMX + Forskolin) with data presented as means ± standard deviations (n=4, MK-571; n=5, Ceefourin-1).

### Detection of ABCC4 and CFTR in human airway epithelial cell lines

The expression and function of ABCC4 has been confirmed in HBEC6-KT cells (**Fig. 1**) although CFTR expression and function remain to be defined^[41,42]^. In contrast, human airway epithelial Calu-3 cells are well documented for CFTR expression and function but not for ABCC4^[53–55]^.

ABCC4 and CFTR protein expression levels were therefore probed via immunoblot in HBEC6-KT and Calu-3 cell lines with total protein stain performed as a loading control (**Fig. 2**). Due to reported time-dependent differentiation and polarization of Calu-3 cells, protein was extracted at 4 different time points (0, 7, 14, and 21 days post-confluency)^[53,56–58]^. ABCC4 was present in both HBEC6-KT and Calu-3 cells at all-time points, indicated by the bands present at 150 kDa. CFTR was present at all-time points in Calu-3 cells. No CFTR was detected in HBEC6-KT cells. Due to co-expression of ABCC4 and CFTR, Calu-3 cells were used for subsequent studies exploring the interrelationships between ABCC4, cAMP modulation, and CFTR.

**Figure 2:**
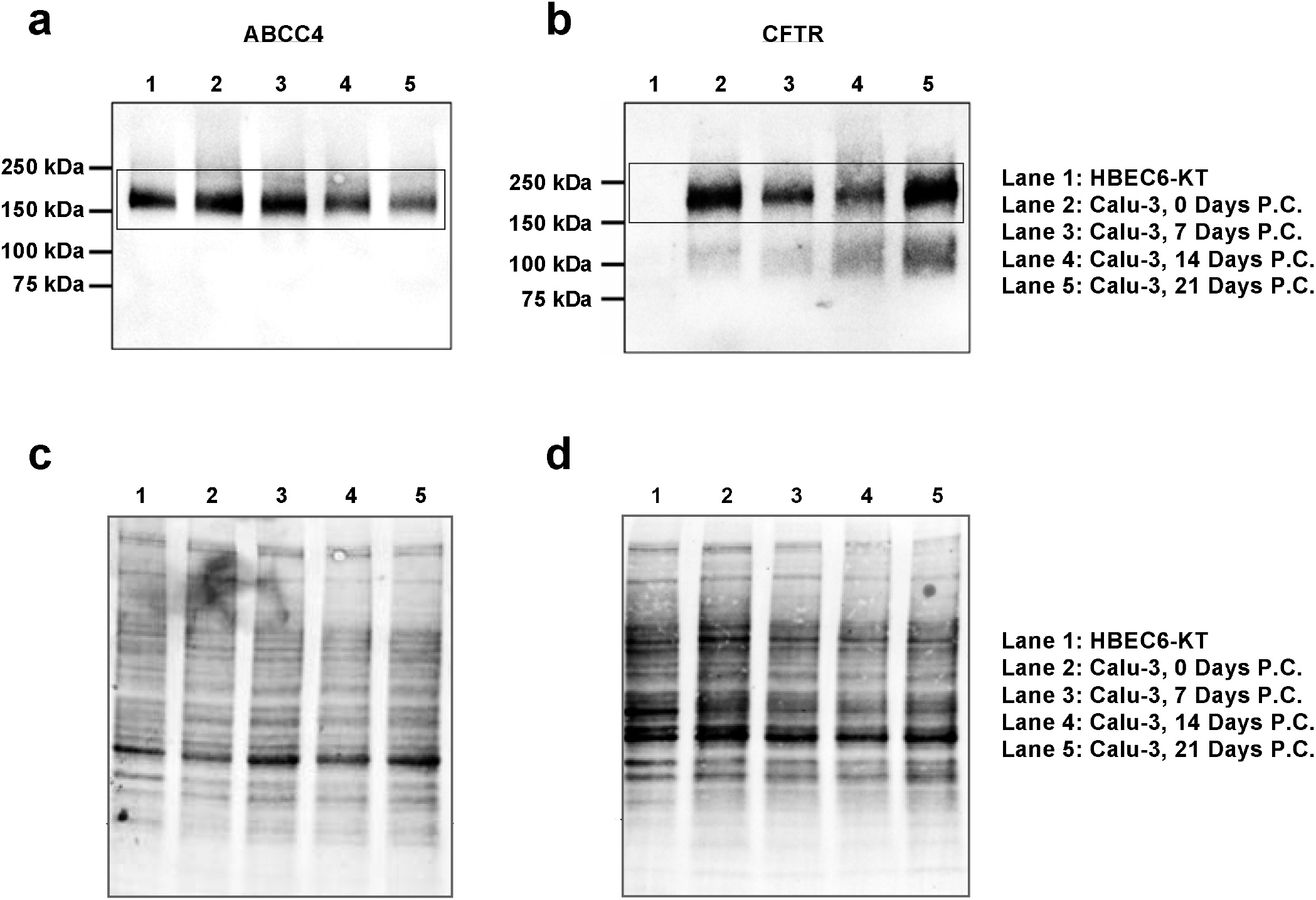
Characterization of ABCC4 and CFTR protein expression in two human airway epithelial cell lines. (**a**) ABCC4 and (**b**) CFTR in both HBEC6-KT and Calu-3 cells. For Calu-3 cells, extracts were retrieved at several different time points post-confluence (P.C.). (**a**) ABCC4 was detected in both HBEC6-KT and Calu-3 cells and (**b**) CFTR was only detected in Calu-3 cells. A total protein stain was performed as the loading control for (**c**) ABCC4 and (**d**) CFTR blots. Full-length blots are presented in Supplementary Fig. 1.

### Receptor-dependent and -independent activation of CFTR activity – Impact of VX-770

A concentration-response analysis was performed with forskolin (FSK) and isoproterenol (ISO), G protein-coupled receptor-independent and dependent cAMP-inducers, respectively (**Fig. 3**). The half maximal effective concentration (EC_50_) value for FSK and ISO were determined to be 0.19 μM and 0.07 μM respectively, with both compounds demonstrating ability to induce CFTR activity as measured by a validated membrane potential sensitive assay^[14]^.

**Figure 3:**
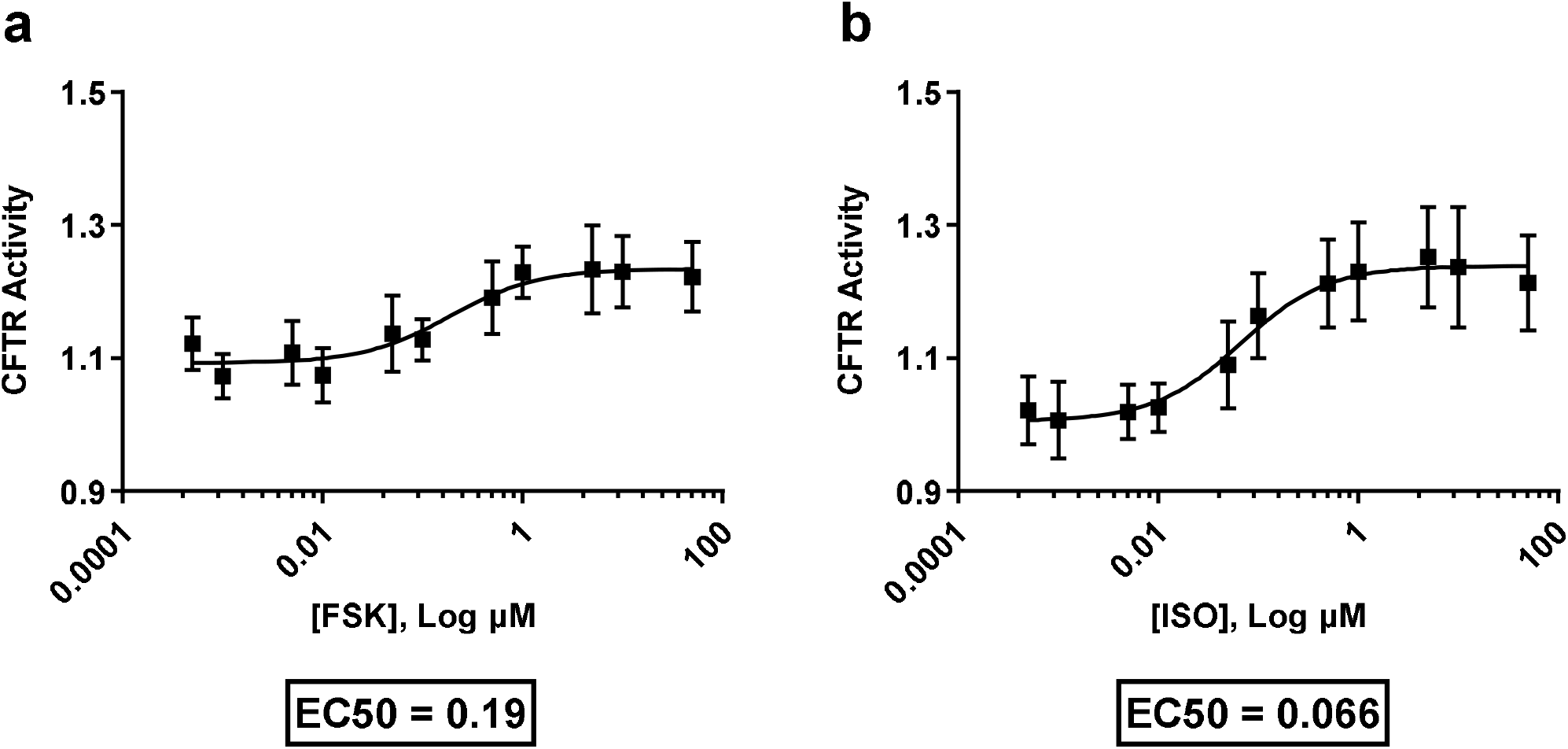
Concentration-response analysis of two different cAMP-elevating agents on CFTR activity. Calu-3 cells were exposed to G protein-coupled receptor-independent cAMP inducer (**a**) forskolin and G protein-coupled receptor-dependent cAMP inducer (**b**) isoproterenol at increasing concentrations while measuring CFTR activity using an *in vitro* CFTR membrane potential assay. EC_50_ values were analyzed and reported. Each concentration-response curve was normalized to baseline over DMSO vehicle control with data presented as ± standard deviations (n=8).

Concentration-response curves with CFTR modulator VX-770, using both receptor-independent FSK and receptor-dependent ISO cAMP inducers (**Fig. 4a and e**), demonstrated an upward and leftward shift in the curve, indicating an increase in CFTR activity. Area under the curve (AUC) analysis showed a significant increase with the addition of VX-770 with either cAMP inducer (**Fig. 4b and f**; *\*\*P≤0.01* and *\*\*\*\*P≤0.0001*). VX-770 increased the sensitivity of FSK-induced CFTR activity as measured by EC_50_ analysis (**Fig. 4c**; \*\**P≤0.01*). However, the maximal effect concentration (E_max_) remained unchanged (**Fig. 4d**). Despite changes in AUC, no changes in EC_50_ or E_max_ were observed with the combination of VX-770 and ISO (**Fig. 4g and h**).

**Figure 4:**
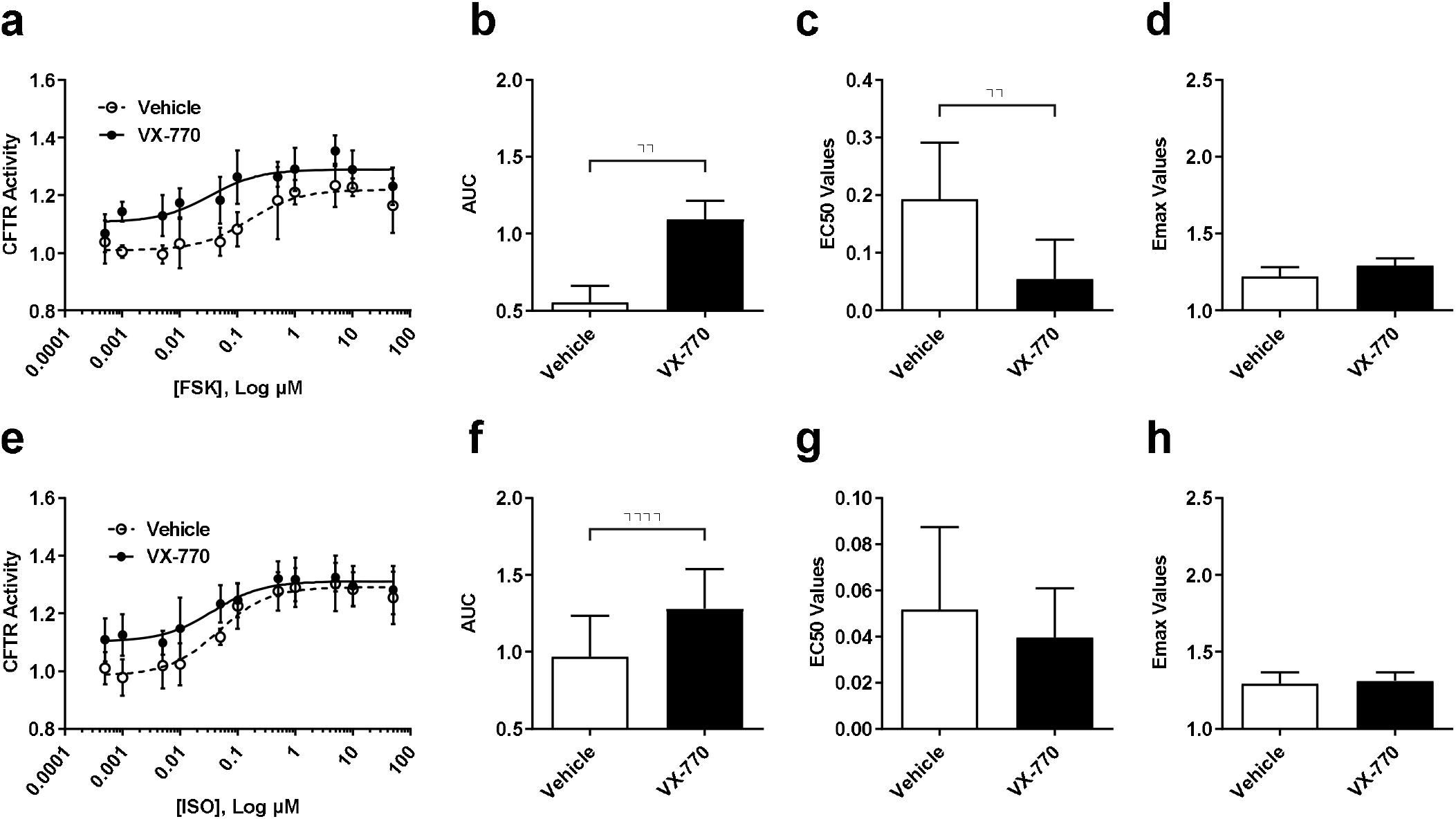
Consequences of CFTR modulator VX-770 on CFTR function using receptor-independent (forskolin) and receptor-dependent (isoproterenol) cAMP inducers. Calu-3 cells were stimulated with VX-770 (1 μM) in the presence of cAMP inducer (**a**) forskolin and (**e**) isoproterenol. Analysis of concentration-response curves was performed for (**b** and **f**) AUC, (**c** and **g**) EC_50_, and (**d** and **h**) E_max_. The concentration-response curve was normalized to baseline over DMSO vehicle control. All data presented as ± standard deviations (n=4, FSK; n=8, ISO). *\*P0.01; ****P≤0.0001*

Collectively, these results demonstrate that both receptor dependent and independent mechanisms of cAMP elevation are able to induce CFTR activity in Calu-3 cells, which in turn is potentiated with VX-770.

### Effect of ABCC4 and PDE-4 inhibition on CFTR activity

Following our quantification of CFTR activity with the clinically approved CFTR potentiator, VX-770, we next investigated the effect of ABCC4 inhibition and PDE-4 inhibition in Calu-3 cells.

For pharmacological inhibition of ABCC4, we used MK-571 and Ceefourin-1 (**see Fig. 1**) with FSK and ISO (**Fig. 5a-d**). No combination of ABCC4 inhibitor and cAMP elevating agent changed CFTR activity in Calu-3 cells when examining AUC, EC_50_, E_max_, and baseline values (**Fig. 5e-h**).

**Figure 5:**
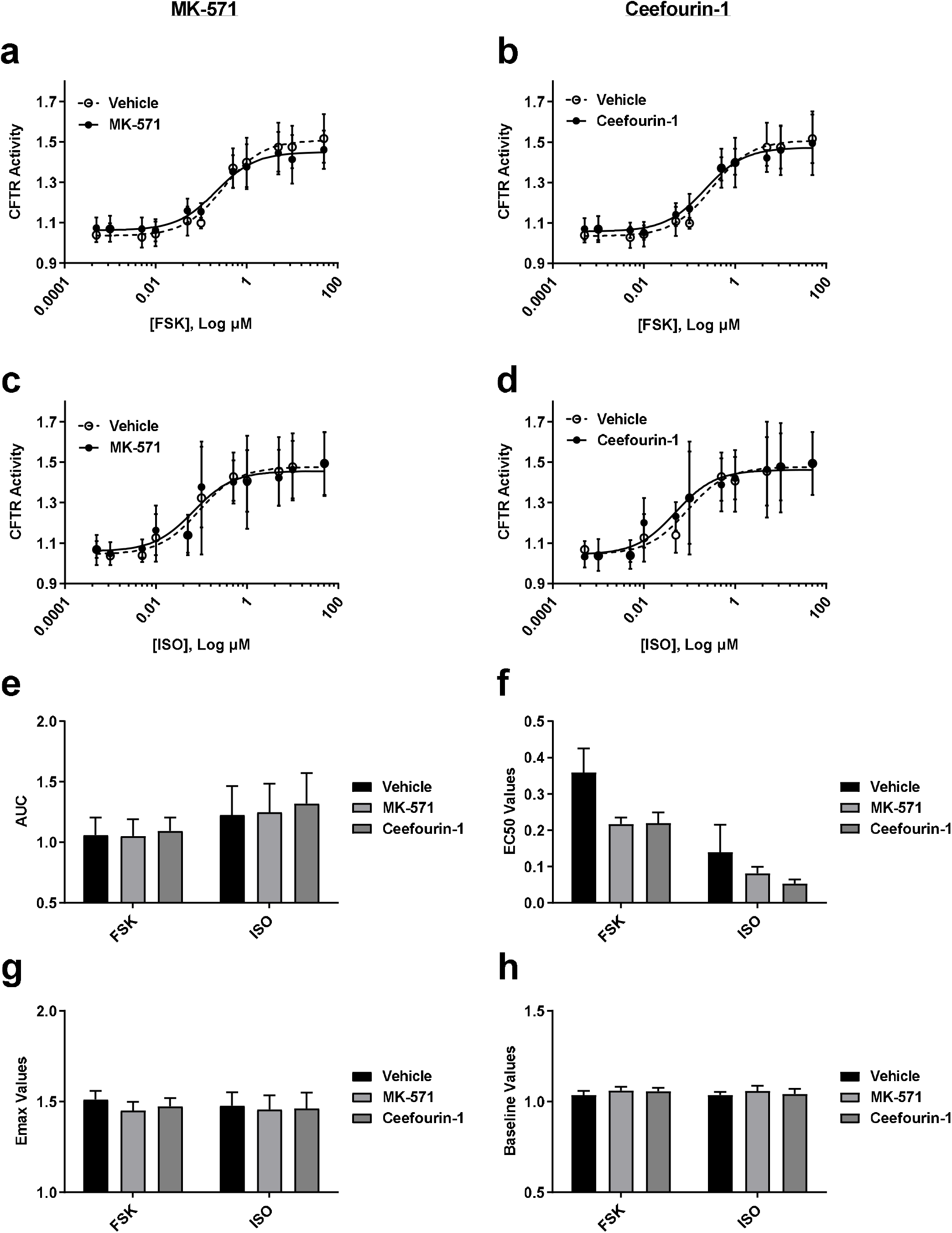
Consequences of ABCC4 inhibition on CFTR function using receptor-independent (forskolin) and receptor-dependent (isoproterenol) cAMP inducers. ABCC4 was inhibited with MK-571 and Ceefourin-1 using submaximal concentrations determined from cAMP-efflux assays. Forskolin stimulated (**a-b**) CFTR activity in the presence of (**a**) MK-571 (1.8 μM) and (**b**) Ceefourin-1 (4.8 μM). Isoproterenol stimulated (**c-d**) CFTR activity in the presence of (**c**) MK-571 and (**d**) Ceefourin-1. Analysis of concentration-response curves was performed for (**e**) AUC, (**f**) EC_50_, (**g**) Emax, and (**h**) Baseline. Each concentration-response curve was normalized to baseline over DMSO vehicle control. All data presented as means ± standard deviations (n=5, FSK; n=4, ISO).

For pharmacological inhibition of PDE-4 we used Roflumilast and Rolipram with FSK and ISO (**Fig. 6a-d**). In contrast to ABCC4 inhibition, the concentration-response curves of PDE-4 inhibitors showed leftward shifts with ISO (**Fig. 6c and d**). Similarly, AUC analysis revealed significant increases with the addition of either Roflumilast or Rolipram with ISO but not FSK (**Fig. 6e**; \**P≤0.05*). EC_50_, E_max_, and baseline values for PDE-4 inhibitors with either cAMP inducer were not different (**Fig. 6f-h**).

**Figure 6:**
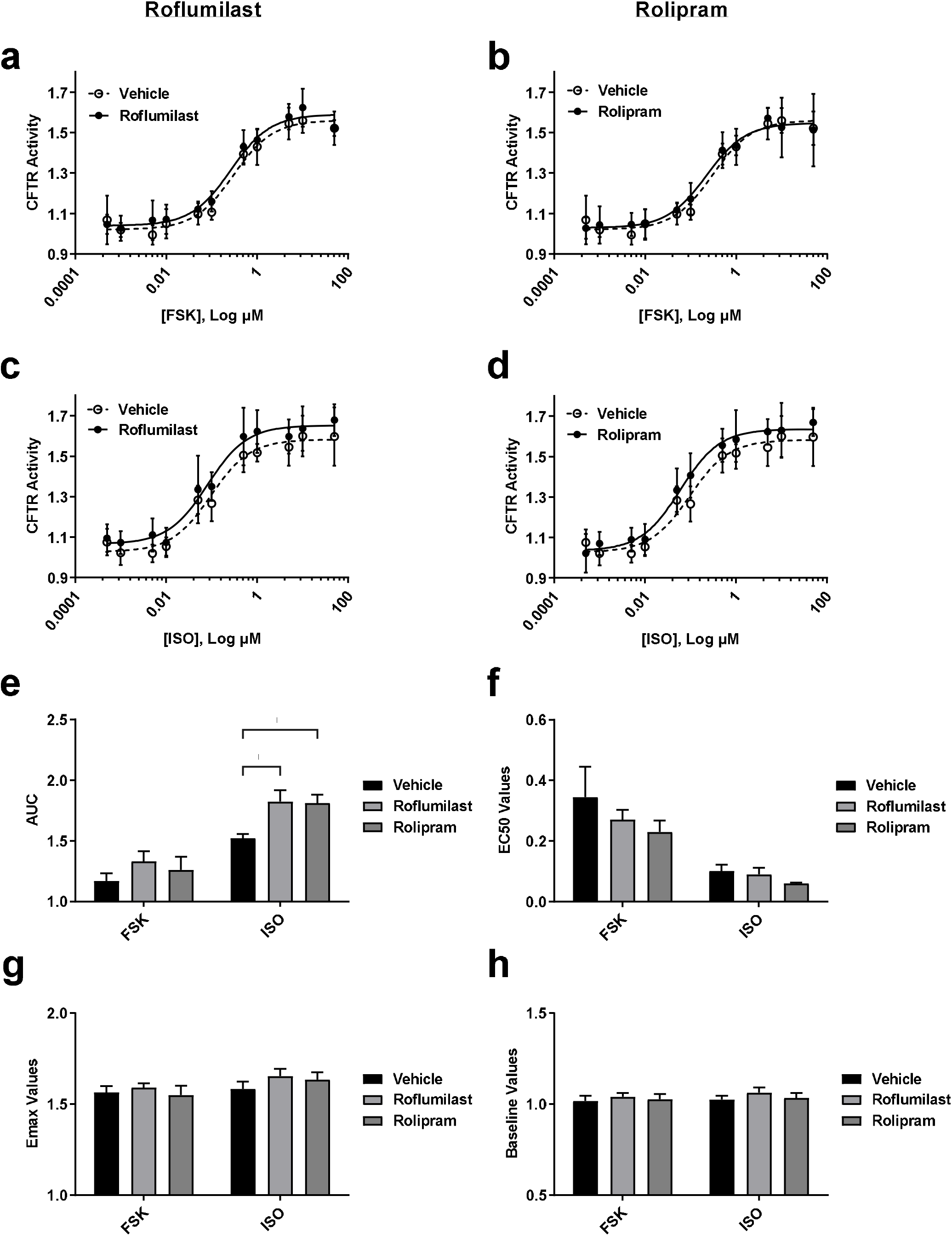
Consequences of PDE-4 inhibition on CFTR function using receptorindependent (forskolin) and receptor-dependent (isoproterenol) cAMP inducers. PDE-4 was inhibited with Roflumilast and Rolipram at submaximal concentrations. Forskolin stimulated (**a-b**) CFTR activity in the presence of (**a**) Roflumilast (1 μM) and (**b**) Rolipram (10 μM). Isoproterenol stimulated (**c-d**) CFTR activity in the presence of (**c**) Roflumilast and (**d**) Rolipram. Analysis of concentration-response curves was performed for (**e**) AUC, (**f**) EC_50_, (**g**) E_max_, and (**h**) Baseline. Each concentration-response curve was normalized to baseline over DMSO vehicle control. All data presented as means ± standard deviations (n=5). \**P*≤0.05

### Effect of cAMP modulation with VX-770 on CFTR activity

Due to our observations that ABCC4 and PDE-4 inhibition alone showed minor potentiation of CFTR activity relative to VX-770, we next investigated whether a combinatorial approach could be more efficacious^[13]^.

Concentration-response curves of VX-770 + MK-571 or Roflumilast combinations with FSK stimulation led to an upward shift of the curve at all concentrations, indicating an increase in CFTR activity (**Fig. 7a and b**). VX-770 + MK-571 or Roflumilast combinations with ISO also led to shifts in the curves, however only at lower concentrations (**Fig. 7c and d**). For the AUC analysis of each combination, increases were observed with FSK but not ISO (**Fig. 7e**; *\*\*P≤0.01*, and *\*\*\*P≤0.001*). EC_50_ and E_max_ analysis for each combination with both cAMP inducers showed no difference (**Fig. 7f and g**).

**Figure 7:**
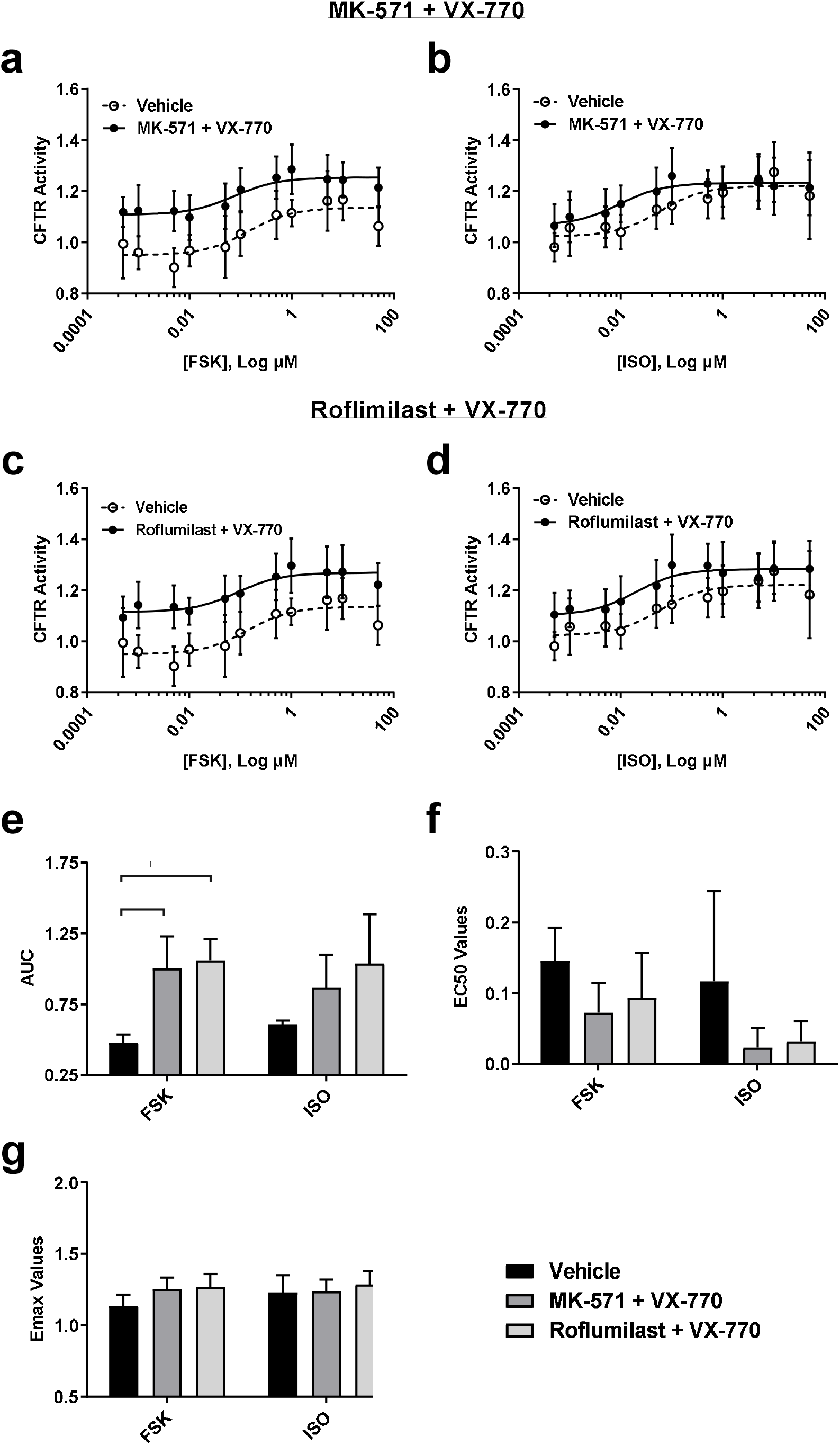
Consequences of ABCC4 and PDE-4 inhibition on CFTR function in the presence of CFTR modulator VX-770 using receptor-independent (forskolin) and receptor-dependent (isoproterenol) cAMP inducers. ABCC4 and PDE4 were inhibited with MK-571 (1.8 μM) and Roflumilast (1 μM) respectively prior to VX-770 (1 μM) stimulation. Forskolin stimulated (**a-b**) CFTR activity in the presence of (**a**) MK-571 + VX-770 and (**b**) Roflumilast + VX-770. Isoproterenol stimulated (**c-d**) CFTR activity in the presence of (**c**) MK-571 + VX-770 and (**d**) Roflumilast + VX-770. Analysis of concentration-response curves was performed for (**e**) AUC, (**f**) EC_50_, and (**g**) E_m_ax. Each concentration-response curve was normalized to baseline over DMSO vehicle control. All data presented as means ± standard deviations (n=6-7, MK-571 + VX-770 and Roflumilast + VX-770). *\*\*P≤0.01;* \*\*\**P*≤0.001

To compare our series of cAMP modulation interventions with and without VX-770 (**Fig. 4-7**), fold-change comparisons of AUC, EC_50_, and E_max_, relative to control conditions were performed for all experiments (**Fig. 8**). Fold-change analysis of AUC with FSK stimulation showed that VX770 was superior to either ABCC4 (MK-571) or PDE-4 (Roflumilast) inhibition alone, and that combinatorial approaches did not potentiate CFTR AUC values (**Fig. 8a**, *\*\*P≤0.01* and *\*\*\*P<0.001*). ISO stimulation resulted in similar modest trends that were not significant (**Fig. 8d**). Fold-change EC_50_ analysis between VX-770 alone to VX-770 with ABCC4 or PDE-4 inhibition showed a significant decrease with ISO for both combinatorial approaches, observations that were absent with FSK stimulation. (**Fig. 8b and e**, \**P≤0.05*). Lastly, fold-change E_max_ analysis performed with either FSK or ISO showed no changes for any combinatorial approach (**Fig. 8c and f**).

**Figure 8:**
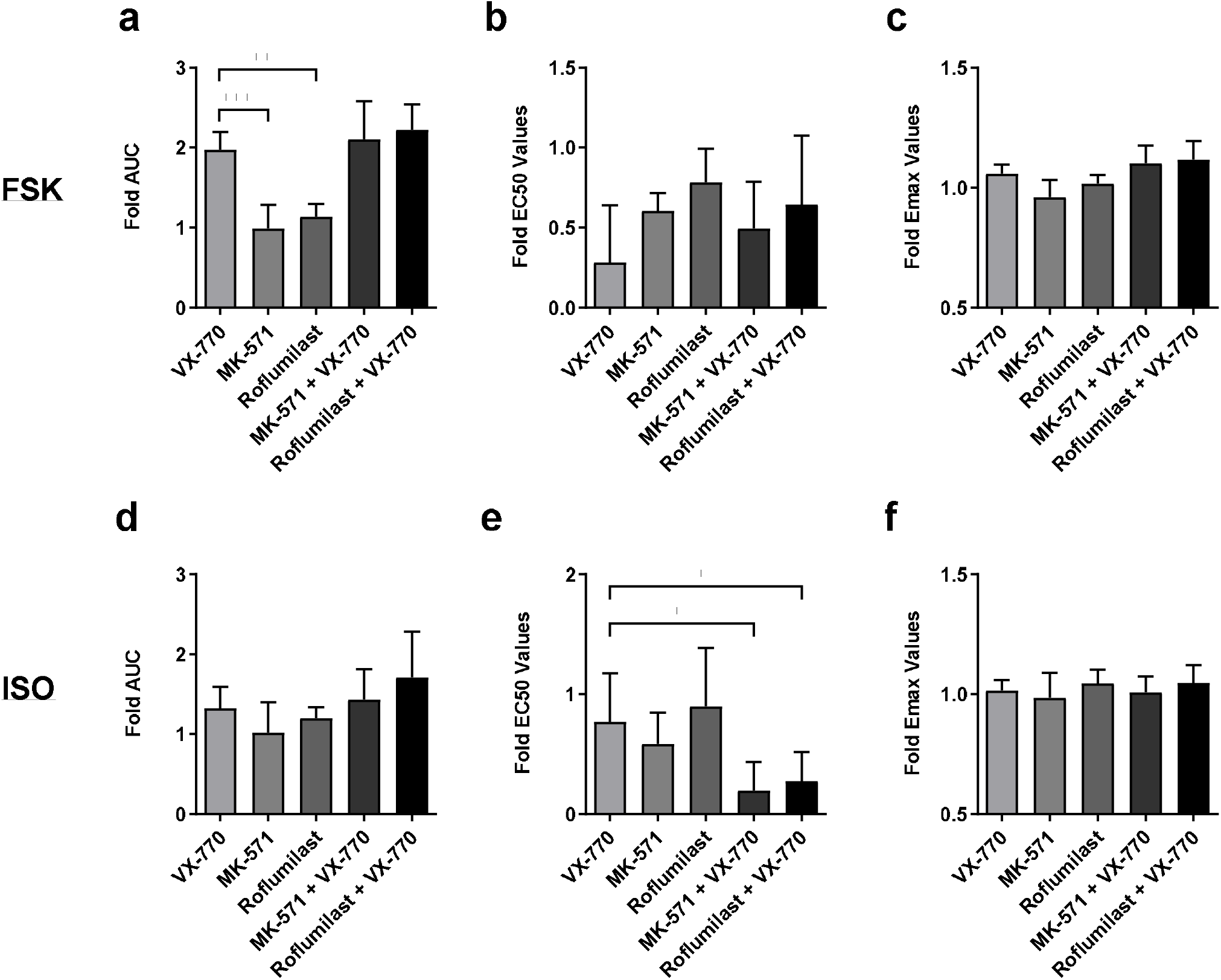
Fold-change comparisons between CFTR modulator VX-770, ABCC4 inhibition, and PDE-4 inhibition. A comparison between CFTR modulator VX-770, ABCC4 inhibitor MK-571, PDE-4 Inhibitor Roflumilast, and VX-770 ± MK-571 or Roflumilast was performed using the values determined from their concentration-response curves in the presence of forskolin (**a-c**) or isoproterenol (**d-f**). Each treatment was normalized to its vehicle control to allow for comparison. A fold-change of (**a** and **d**) AUC, (**b** and **e**) EC_50_, and (**c** and **f**) E_max_ are depicted. All data presented as means ± standard deviations (n=4-8, VX-770; n=4-5, MK-571; n=5, Roflumilast; n=6-7, MK-571 + VX-770 and Roflumilast + VX-770). \**P*≤0.05; *\*\*P≤0.01;* \*\*\**P*≤0.001

## Discussion

There is an increased need for combinatorial therapies for CF subjects to optimally treat all subjects at a personalized level^[13,25,27,29]^. CFTR potentiation with VX-770 represents a high-water mark for increased chloride conductance in CF patients. In the present study, we quantified the capacity of combinatorial approaches that modulate intracellular cAMP levels to activate CFTR. Specifically, we inhibited ABCC4, a cAMP efflux pathway, and PDE-4, a cAMP metabolism pathway. Using Calu-3 cells, we demonstrate that interventions targeting intracellular cAMP are unable to potentiate CFTR to levels observed with VX-770. Combinatorial approaches of cAMP modulation with VX-770 suggest that increases in CFTR activity as measured by AUC and EC_50_ values, are possible and warrant further exploration in primary human airway epithelial cells expressing loss of function CFTR variants^[14]^.

The addition of VX-770 to Calu-3 cells induced shifts in the curve with both FSK and ISO, providing context for the amount of CFTR potentiation possible in Calu-3 cells with a clinically approved CFTR potentiator. While there was a significant change to EC_50_ with FSK, no changes were seen with iSo, suggesting that mechanisms governing cAMP elevations may influence downstream analyses of CFTR activities^[59–63]^. For both cAMP inducers, the E_max_ values were not significant and while there were no significant changes to baseline with FSK, there was a significant increase in baseline with ISO (data not shown). Collectively, this suggests not only is the mechanism of action in elevating cAMP levels affecting sensitivity, but also efficacy^[59–63]^. Collectively, these results suggest that it is possible to potentiate CFTR activity in Calu-3 cells by modulating cAMP levels.

Combinatorial approaches have been suggested as the future for modulation of CFTR activity and may include direct or indirect approaches^[13]^. Modulation of cAMP levels functions as an indirect mechanism to modulate CFTR and could potentially be combined with direct CFTR modulator approaches. We therefore decided to investigate ABCC4 and PDE-4 inhibitors which are able to differentially modulate cAMP levels by blocking extracellular transport and intracellular breakdown, respectively.

It has been previously demonstrated that pharmacological inhibition of ABCC4 by MK-571, in combination with VX-770, was able to improve CFTR activity in primary nasal epithelial cells from CF patients heterozygous for the G551D-variant^[14]^. In addition, in primary non-CF human bronchial epithelial (HBE), it has been demonstrated that the administration of Roflumilast led to CFTR activation^[38]^. However, in this same study, the addition of Roflumilast to Calu-3 cells did not potentiate CFTR activity beyond the max stimulation that resulted from the addition of FSK^[38]^.

In our concentration-response analysis of ABCC4 and PDE-4 inhibitors in Calu-3 cells, only mild CFTR potentiation was observed alone or in combination with VX-770. Our results for ABCC4 inhibition contrast our previous demonstration of ABCC4 inhibitor MK-571 in potentiating CFTR activity in G551D-variant expressing primary human airway epithelial cells^[14]^. Conversely, our results of PDE-4 inhibitors align with previous findings in primary HBE cells that were exposed to whole cigarette smoke in which Roflumilast, in combination with VX-770, demonstrated an increase in short-circuit current, suggesting this combination leads to an increase in CFTR activity^[38]^. We highlight that it may be possible that wild-type CFTR expressed in Calu-3 cells is maximally active and difficult to further potentiate with cAMP modulating agents. Our data showing shifts in the left of the curve (increased sensitivity of CFTR activity) without changes in maximal response support this concept. Specifically, the addition of VX-770 with either cAMP inducer to Calu-3 cells led to an upward shift in the curve, a significant increase in AUC but no significant changes to E_max_. However, VX-770 with FSK showed a significant decrease in EC_50_ and VX-770 with ISO showed a significant increase in baseline values. When VX-770 was combined with an ABCC4 or PDE-4 inhibitor and cAMP inducer FSK, there were significant increases in AUC, indicating an increase in CFTR activity, but again no change in E_max_. In order to determine whether ABCC4 and PDE-4 inhibitor with VX-770 had an additive effect beyond VX-770 alone, a fold-change analysis for AUC, EC_50_, and E_max_ values of the independent experiments was performed for indirect comparisons. Combinatorial additions of VX-770 with ABCC4 or PDE-4 inhibitor, in the presence of cAMP inducer ISO, led to a decrease in fold EC_50_ values compared to VX-770 alone, suggesting that ABCC4 and PDE-4 inhibitors may increase the sensitivity to VX-770. In addition, with cAMP inducer FSK, VX-770 with Roflumilast had a significant increase in fold baseline values, suggesting that there may also be an increase in efficacy. With these fold-change comparison analyses in mind, further investigation of these combinatorial treatments should be performed.

Our interrogation of the role of ABCC4 in airway epithelial cell modulation of cAMP levels is grounded in reports from primary human airway epithelial cells and cell lines, while examining CFTR activity in Calu-3 cells has a strong foundation^[35,41,42,53,54]^. To determine the optimal cell system to perform ABCC4 interventions with outcomes of CFTR activity, we interrogated both HBEC-6KT and Calu-3 cells for ABCC4 and CFTR protein expression levels. To our surprise, while ABCC4 is present in both cell lines, CFTR is only present in the Calu-3 cells, leading to the selection of Calu-3 cells as the model of study. Our observation highlights the importance of basal expression analysis for CFTR protein in any human airway epithelial cells or lines prior to performing functional experiments.

Elevations in intracellular cAMP can be induced via multiple G protein-coupled receptordependent (ß-adrenergic receptor and adenosine receptor agonists - ISO and adenosine respectively) and independent mechanisms (direct activation of the enzyme adenylyl cyclase - FSK)^[64-68]^. The compartmentalization of cAMP signalling may be important for downstream signalling pathways within the cell - with specific implications for CFTR activity^[34,69]^. To determine whether the mechanism of cAMP production impacts CFTR activity, Calu-3 cells were treated with FSK and ISO at varying concentrations. Sigmoidal shape concentration-response curves were observed for both cAMP elevating agents with the maximal degree of change for CFTR activity greater with ISO. The interpretation of this observation is that the compartmentalization of cAMP to the plasma membrane may more effectively couple protein kinase A activation to CFTR phosphorylation, which could be masked by global cytosolic increases in cAMP induced by FSK^[34,69]^.

In our previous studies, we have not performed complete concentration-response analysis experiments for the leukotriene receptor antagonist MK-571, which has off-target ABCC4 inhibition effects, in parallel to the more selective ABCC4 inhibitor Ceefourin-1, thus we wanted to explore both in greater detail^[14,42,45]^. In our concentration-response analysis experiments utilizing an *in vitro* cAMP-efflux assay platform, both commercially available Ceefourin-1 and MK-571 were able to attenuate extracellular transport of cAMP, thus both were used in downstream *in vitro* CFTR membrane potential assays to investigate the consequences of ABCC4 inhibition on CFTR function. Due to similar findings using either compound in the *in vitro* CFTR membrane potential assay and previous demonstrations of MK-571, in combination with VX-770, potentiating CFTR activity beyond VX-770 alone, MK-571 was selected for combinational studies^[14]^.

Further investigation of ABCC4 and PDE-4 inhibitors for combinatorial therapy should be performed in human airway epithelial cells from CF subjects with compromised CFTR expression or function. Although Calu-3 cells expressed both ABCC4 and CFTR, it might not have been an appropriate model of study due to its WT-CFTR expression. The potentiation of CFTR activity by ABCC4 or PDE-4 inhibitors in the literature were performed in primary cells, suggesting that a potential add-on combinatorial therapy may only show an effect in primary cell cultures, especially ones with CFTR defects.

## Methods

### Reagents

The human bronchial epithelial cell (HBEC6-KT) line was cultured in Keratinocyte Serum-Free Medium 1X (Gibco^®^) according to the manufacturers directions and supplemented with P/S (VWR^®^) at 1X^[70]^. Human airway epithelial cell line Calu-3 (ATCC HTB-55™) were cultured in Minimum Essential Medium Alpha Medium (Corning^®^) and supplemented with Premium Grade FBS (VWR^®^) at 10%, HEPES (Corning^®^) at 1X, and P/S (VWR^®^) at 1X. The following chemical reagents were dissolved in DMSO, cAMP elevating agents Forskolin (Cayman Chemical) and Isoproterenol (Cayman Chemical), ABCC4 inhibitors MK-571 (Cayman Chemical) and Ceefourin™ 1 (Abcam), PDE inhibitors Roflumilast (Cayman Chemical) and Rolipram (Cayman Chemical), and CFTR Inhibitor (Selleck Chemicals).

### Human airway epithelial cell lines

Two human airway epithelial cell lines were used for *in vitro* experiments. HBEC6-KT cells, derived from healthy non-smoking individuals and immortalized through human telomerase reverse transcriptase and cyclin-dependent kinase 4 expression was used for *in vitro* concentration-response analysis of ABCC4 inhibitors through a cAMP-efflux assay^[41,42,70–72]^. Calu-3 cells derived from the metastatic site of lung adenocarcinoma tissue was used for *in vitro* analysis of CFTR function.

### ABCC4 and CFTR protein expression analysis

HBEC6-KT and Calu-3 cells were lysed using RIPA Lysis Buffer containing Protease Inhibitor Cocktail powder (Sigma-Aldrich) for 60-90 minutes at 4°C on a rocker. Lysates were then centrifuged at 16,000xg for 15 minutes with the supernatants collected for downstream immunoblots for ABCC4 and CFTR. Protein quantification was performed using a BCA protein assay. Twenty micrograms of protein were loaded on 4-15% gradient TGX Stain-Free™ Protein Gels and transferred to Immun-Blot^®^ LF PVDF membrane (Bio-Rad). The membranes were blocked with 1X TBS with 0.05% Tween 20 and 5% skim milk powder for 2 hours at 25°C. The membranes were then incubated with primary antibody ABCC4/MRP4 (1:40, Abcam, AB15602) or CFTR (1:5000, UNC-Chapel Hill, AB596) overnight. The membranes were then washed in 1X TBS with 0.05% Tween 20 and incubated with HRP-linked anti-rat secondary antibody (1:3000, Cell Signaling Technology^®^, 7077S) or anti-mouse secondary antibody (1:3000, Cell Signaling Technology^®^, 7076S) for 2 hours at 25°C. A chemiluminescence image of the blot was taken using the Bio-Rad Image Lab software.

### *In vitro* cAMP-efflux assay

HBEC6-KT cells were cultured as described previously^[41,42,71,72]^. Cells were pretreated with IBMX (20 μM) for 2 hours prior to experimental conditions. After the pre-treatment of IBMX, cells were treated with ABCC4 inhibitors (0.01, 0.1, 1, 10, or 100 μM) or DMSO for 30 minutes. Following these exposures, the cells were treated with forskolin (10 μM) for 6 hours and cellculture supernatants were collected for analysis of extracellular cAMP. The negative control was IBMX alone and the positive control was IBMX and forskolin.

### *In vitro CFTR* membrane potential assay

An *in vitro* CFTR membrane potential assay was performed on Calu-3 cells cultured in 37°C as previously described^[14]^. Calu-3 cells were grown for 21 days on 96-well plates and washed with HBSS prior to experimental conditions. After washing with HBSS, cells were loaded with BLUE Membrane Potential Dye (Molecular Devices, #R8042) dissolved in 37°C chloride-free buffer (NMDG Gluconate Buffer - 150 mM NMDG-gluconate, 3 mM KCl, 10mM HEPES, pH 7.35, 300 mOsm). Immediately after dye loading, CFTR activity measurements were completed during baseline (40 minutes), cAMP elevation (30 minutes), and CFTR inhibition (20 minutes). For assays with ABCC4 or PDE-4 inhibitors, a pre-incubation with the inhibitors (30 minutes) in the plate reader was performed prior to cAMP elevation. CFTR modulator VX-770 was used at 1 μM. cAMP-elevation was performed using forskolin or isoproterenol (0.0005-50 μM). CFTR inhibition was performed with CFTRinh-172 (10 μM). At the end of the assay, raw data was exported for statistical analysis. CFTR activity is determined by dividing the membrane potential peak after the ABCC4 or PDE-4 inhibition and cAMP elevation additions to the stabilized baseline over DMSO control.

### Statistical analysis

For the *in vitro* cAMP-efflux assay, values were normalized to the positive control (IBMX and forskolin) and the half maximal inhibitory concentration (IC_50_) was determined. SD was calculated using data from biological replicates (n=4-5). For *in vitro* CFTR membrane potential assays, values were normalized to the averaged baseline over averaged vehicle (DMSO) control of all biological replicates. SD was calculated using data from biological replicates (n=4-8). An area under the curve (AUC), half-maximal concentration (EC_50_), maximum concentration (E_max_), and baseline analysis was performed for each individual replicate then averaged together. The AUC analysis is an aggregate measure of sensitivity and maximal response, allowing for the observation of the total net responsiveness of the cells. The AUC analysis was done by normalizing the individual replicates to the bottom most value of the averaged vehicle control. In order to compare the independent experiments performed, a fold-change analysis using individual AUC, EC_50_, and E_max_, determined from their respective concentration-response curves, was performed and was normalized to the averaged DMSO vehicle control. A one-way ANOVA with subsequent post-hoc test or a paired *t*-test was performed. Statistical analysis was performed using GraphPad Prism 6.

### Data Availability Statement

The datasets generated and analysed during the course of the study are available from the corresponding author on reasonable request.

## Acknowledgements

This research was supported by funding from the Canadian Institutes of Health Research (CIHR).

## Author Contributions

J.P.N. performed *in vitro* CFTR membrane potential assay experiments, data analyses, figure generation, and literature review; contributed to the manuscript conception; and drafted the manuscript. M.B. performed *in vitro* CFTR membrane potential assay experiments. R.D.H performed *in vitro* cAMP-efflux assay experiments, data analyses, and figure generation. N.T. performed western blotting and figure generation. M.D.I. contributed to the manuscript conception. J.A.H. (Principle Investigator and corresponding author) was responsible for the oversight of the entire study (data collection, analysis, drafting, finalization) and supervision of trainees.

## Competing Interests

The authors declare no competing interests.

## Notes

### Competing Interest Statement

The authors have declared no competing interest.

